# Keeping it clean: the cell culture quality control experience at the National Center for Advancing Translational Sciences

**DOI:** 10.1101/798629

**Authors:** Jacob S. Roth, Tobie D. Lee, Dorian M. Cheff, Maya L. Gosztyla, Rosita Asawa, Carina Danchik, Sam Michael, Anton Simeonov, Carleen Klumpp-Thomas, Kelli Wilson, Matthew D. Hall

## Abstract

Quality control monitoring of cell lines utilized in biomedical research is of critical importance, and is critical for reproducibility of data. Two key pitfalls in tissue culture are 1) cell line authenticity, and 2) mycoplasma contamination. As a collaborative research institute, the National Center for Advancing Translational Sciences receives cell lines from a range of commercial and academic sources, that are adapted for high-throughput screening. Here, we describe the implementation of a routine NCATS-wide mycoplasma testing and short-tandem repeat (STR) testing for cell lines. Initial testing identified a >10% mycoplasma contamination rate, but the implementation of clearly defined protocols that included immediate destruction of contaminated cell lines wherever possible has led to a much-reduced mycoplasma contamination rate, and data for >2,000 cell line samples tested over 3 years, and case studies are provided. STR testing of 170 cell lines with established STR profiles revealed only 5 mis-identified cell lines received from external labs. The data collected over the three years since implementation of this systematic testing demonstrates the importance of continual vigilance for rapid identification of ‘problem’ cell lines, for ensuring reproducible data in translational science research.

## Introduction

Reproducibility and quality of scientific data are influenced by a wide range of factors, and require clear experimental methodologies to be reported, as well as access to equivalent experimental reagents where possible. Reproducibility also requires characterization of reagents to ensure that they are unsullied. Cell culture (often referred to as tissue culture) is a cornerstone of modern biomedical research. Multiple quality control measures are critical for ensuring that reported data is reproducible, including cell line authentication and confirmation that cell lines are not contaminated.

**Cell line authentication** is essential to ensure that cell lines have not been accidentally mislabeled or cross-contaminated, leading to inaccurate interpretations based on disease models^1,2^. Perhaps the most notorious example of cell line misidentification is that of the HeLa human adenocarcinoma cell line, which has cross-contaminated many other cell lines^3^. The high doubling rate of HeLa cells means that a small number of contaminating cells can outgrow and overtake a cell line it has contaminated. One of the better-known examples of this is the KB cell line. It was derived by Harry Eagle as an epidermoid carcinoma in the 1950s and shown a decade later to be a HeLa contaminant line^4^. In another instance of cell line cross-contamination, MCF7 human breast cancer cells selected for resistance to the chemotherapeutic adriamycin (originally named MCF-7/AdrR) were later found to be derived from the human ovarian carcinoma cell line OVCAR-8. The resistant line was one of the seminal NCI60 cell line panel and has been renamed NCI/ADR-RES^5^. In the instance of the rare cancer adenoid cystic carcinoma, the few cell lines available to the scientific community were shown to be mis-identified, and included rat and mouse cell lines^6^. To avoid future misidentifications, DNA profiling of human cell lines by short tandem repeat (STR) analysis upon receipt, and periodically during culture, can verify the identity of a cell line and mitigate severe and costly repercussions^7,8^.

**Mycoplasma contamination** has similarly plagued the practice of cell culture. Mycoplasma are single-celled prokaryotes that are devoid of a cell wall. Contamination of *in vitro* cultures by *mycoplasma* was first reported in 1956^9^, and has continued to present a challenge to routine culture ever since. The primary source of mycoplasma in continuous cell culture is believed to originate from the human oral cavity (i.e. laboratory cell culture operators, speaking and breathing^10,11^). This hypothesis was supported by one author upon testing and confirming *mycoplasma* presence in their own saliva. More recently, there has been an increasing awareness of the impact of *mycoplasma* contamination on reproducibility in the scientific literature. Screening of cell culture collections have produced estimates that 15-35% of cell lines are *mycoplasma* contaminated^12,13^ (and higher numbers have been reported in individual collections). The recognition of the ease with which tissue culture can be contaminated and the implications for industries such as biotherapeutic production have stimulated the development of off-the-shelf mycoplasma detection kits to allow routine laboratory screening for mycoplasma contamination. Guidelines for systematic testing of cell culture have been previously published^14^.

While *mycoplasma* infection can easily go unnoticed given that infected cells often do not present with visible or morphological symptoms, *mycoplasma* contamination of cell lines can broadly impact cellular biology. These perturbations can alter DNA, RNA, protein synthesis, metabolics, and general cellular processes, though few examples of systematic studies of the impact of mycoplasma have been reported^15^. Microarray analyses on contaminated human cells in culture have reported that *mycoplasma* can affect expression of hundreds of genes, including those encoding ion channels, receptors, growth factors, and oncogenes^16,17^.

A number of phenotypes that are reported to be due to *mycoplasma* contamination can be misinterpreted as impacting the underlying human biology and can produce plausible but irreproducible data that impede translational science. For cancer experimental therapeutics, *mycoplasma* has been shown to affect cancer cell line response to chemotherapy. For example, Liu and colleagues noted increased sensitivity to cisplatin, gemcitabine, and mitoxantrone in HCC97L human hepatocarcinoma cells infected with *Mycoplasma hyorhinis* compared to uninfected cells^18^. *Mycoplasma* contaminated HCT-116 colon cancer cells were found to be 5- to 100-fold more resistant to 5-fluorouracil and 5-fluorodeoxyuridine, respectively, compared to cells that were ‘cured’ of infection with antibiotics^19^. One of the authors has published a report that tiopronin (thiola) selectively kills multidrug-resistant (MDR) cancer cell lines^20^, but subsequent to publication it was found that the MDR cells were mycoplasma contaminated^21^. Treating the cells with plasmocin to remove *mycoplasma* contamination reversed the sensitivity of the cells to tiopronin, and resistant cell lines were sensitized to tiopronin when intentionally contaminated with mycoplasma. As a consequence, the authors elected to retract the publication and correct the record. While the data was legitimate, the scientific conclusions were not valid (‘tiopronin kills mycoplasma-contaminated drug-resistant cell lines’)^21^. For each individual example above, there are likely thousands of unrecognized examples in the literature.

### The mycoplasma testing experience at NCATS

At the NIH National Center for Advancing Translational Sciences (NCATS), there is a strong scientific focus on collaborating with the scientific community. Using assay development and high-throughput screens, and advanced technologies such as induced pluripotent stem cells and tissue printing, we are working to further our understanding of rare and neglected diseases, novel targets, and expand basic biological understanding of the “undrugged” genome. This is accomplished through a team science approach that often includes assay development and automated quantitative high-throughput screening (qHTS) with a small molecule library to identify active hits in biochemical or cell-based assays^22^. Hits may progress to medicinal chemistry to develop a small molecule probe, guided by orthogonal and cell-based counter assays. In such a collaborative environment, NCATS routinely receives cell lines from partnering labs and a range of commercial vendors.

Given the phenotypic impact of mycoplasma contamination, executing a HTS of hundreds of thousands of compounds with such a cell line would be costly and wasteful. Accepting the reality that mycoplasma-contaminated cell lines may regularly be received from collaborators, or that contamination may arise during culture in labs at NCATS, we established a routine weekly mycoplasma testing system. A central location was established for NCATS scientists to deliver a sample of expended culture media, a ‘MycoAlert’ (Lonza) assay was adapted for 96-well plates (scaling the volume by one-half), and samples were tested each Friday with results emailed to those submitting samples. The MycoAlert assay couples the production of ATP (from a provided substrate) by an endogenous Mycoplasma enzyme, with a luciferase enzyme to produce light (chemiluminescence). If the ratio of luminescence from post to pre-readings is >1.2, the sample is positive for Mycoplasma (<0.9 is negative and 0.9 ≤ 1.2 indicates an ambiguous result). Implementation was achieved with minimal burden: the assay is affordable and can be accomplished by a post-baccalaureate fellow in the laboratory within one hour. A weakness of the MycoAlert assay is that it can only detect a limited number of *Mycoplasma* species, while PCR-based methods of mycoplasma detection are highly sensitive due to species specific primers. However PCR-based detection are more time consuming and costly than MycoAlert.

A set of policies were implemented for use of cell lines for discovery purposes in the NCATS biology and high-throughput screening labs in tandem with this weekly testing protocol:

1. All cell lines must be certified mycoplasma-free prior to receipt. As part of establishing a collaboration, any cell lines developed by the collaborating laboratory that are to be shipped to NCATS must be affirmed as mycoplasma-free by the collaborator.
2. All cell lines are tested at NCATS upon receipt. When cell lines received are thawed into culture for the first time, they are tested to confirm they are mycoplasma-free.
3. All cell lines in regular culture must be tested at least once a month, or when thawed from cryovial stocks.
4. All cell lines must be tested immediately prior to HTS execution. As part of planning for an automated HTS, cell culture scale-up must include re-testing for mycoplasma. The negative result must be presented as part of the assay protocol hand-over to the automation team.
5. Contaminated (‘positive’) cell lines should be destroyed immediately, and back-up frozen stock cultured and assessed for mycoplasma status. All stock should be destroyed if other cryovials prove to be positive. In exceptional circumstances, contaminated cells can be quarantined in a dedicated incubator outside the tissue culture room, and plasmocin ‘treatment’ can be used to destroy culture (an example situation is described below).
6. Cell lines that receive ‘ambiguous’ results should be quarantined and re-tested the following week. Two ambiguous tests should be considered contaminated and dealt with as outlined in (5) above.

Since routine mycoplasma testing was implemented by us at NCATS, just over 2,000 samples have been tested. In the three full calendar years of testing to date (2016, 2017, and 2018) an average of 500 samples were tested per year (Table 1). In the first year of testing, 15% of cell line samples were positive (Table 1). A month-by-month breakdown of historical data (Figure 1) shows a 33% positive return rate in the first month. These data are consistent with other reports on contamination rate^12,13^, and perhaps unsurprising given that routine testing had not been implemented previously. Subsequent to the first year of testing, 6-8% of cell lines tested positive (Table 1), with occasional ‘spikes’ (for example, see November 2018 and April 2019, Figure 1) that were associated with testing and re-testing of a number of cell lines received from external laboratories. While cell culture can occasionally suffer from fungal or bacterial contamination across tissue culture flasks that is visually evident in culture medium^23^, our anecdotal evidence is that no such ‘outbreak’ of mycoplasma contamination was observed over the almost four-year period.

**Table 1:**
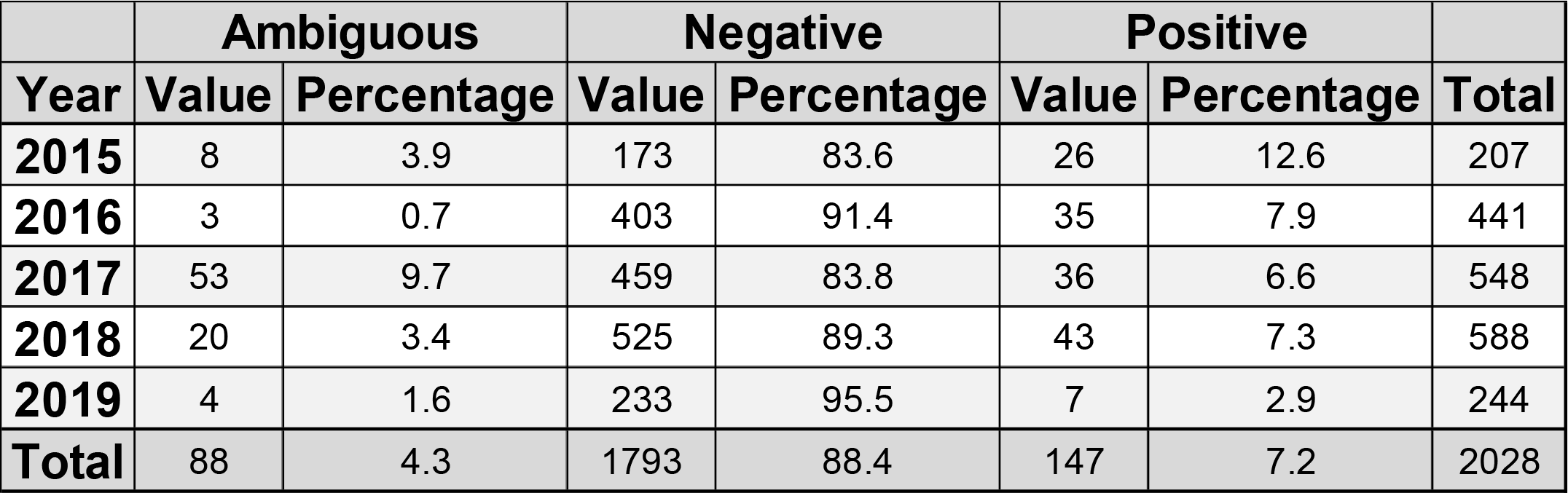
Data are presented as cumulative for each year, with percentages for each outcome.

|  | **Ambiguous** |  | **Negative** |  | **Positive** |  |  |
| --- | --- | --- | --- | --- | --- | --- | --- |
| **Year** | **Value** | **Percentage** | **Value** | **Percentage** | **Value** | **Percentage** | **Total** |
| **2015** | 8 | 3.9 | 173 | 83.6 | 26 | 12.6 | 207 |
| **2016** | 3 | 0.7 | 403 | 91.4 | 35 | 7.9 | 441 |
| **2017** | 53 | 9.7 | 459 | 83.8 | 36 | 6.6 | 548 |
| **2018** | 20 | 3.4 | 525 | 89.3 | 43 | 7.3 | 588 |
| **2019** | 4 | 1.6 | 233 | 95.5 | 7 | 2.9 | 244 |
| **Total** | 88 | 4.3 | 1793 | 88.4 | 147 | 7.2 | 2028 |

**Figure 1:**
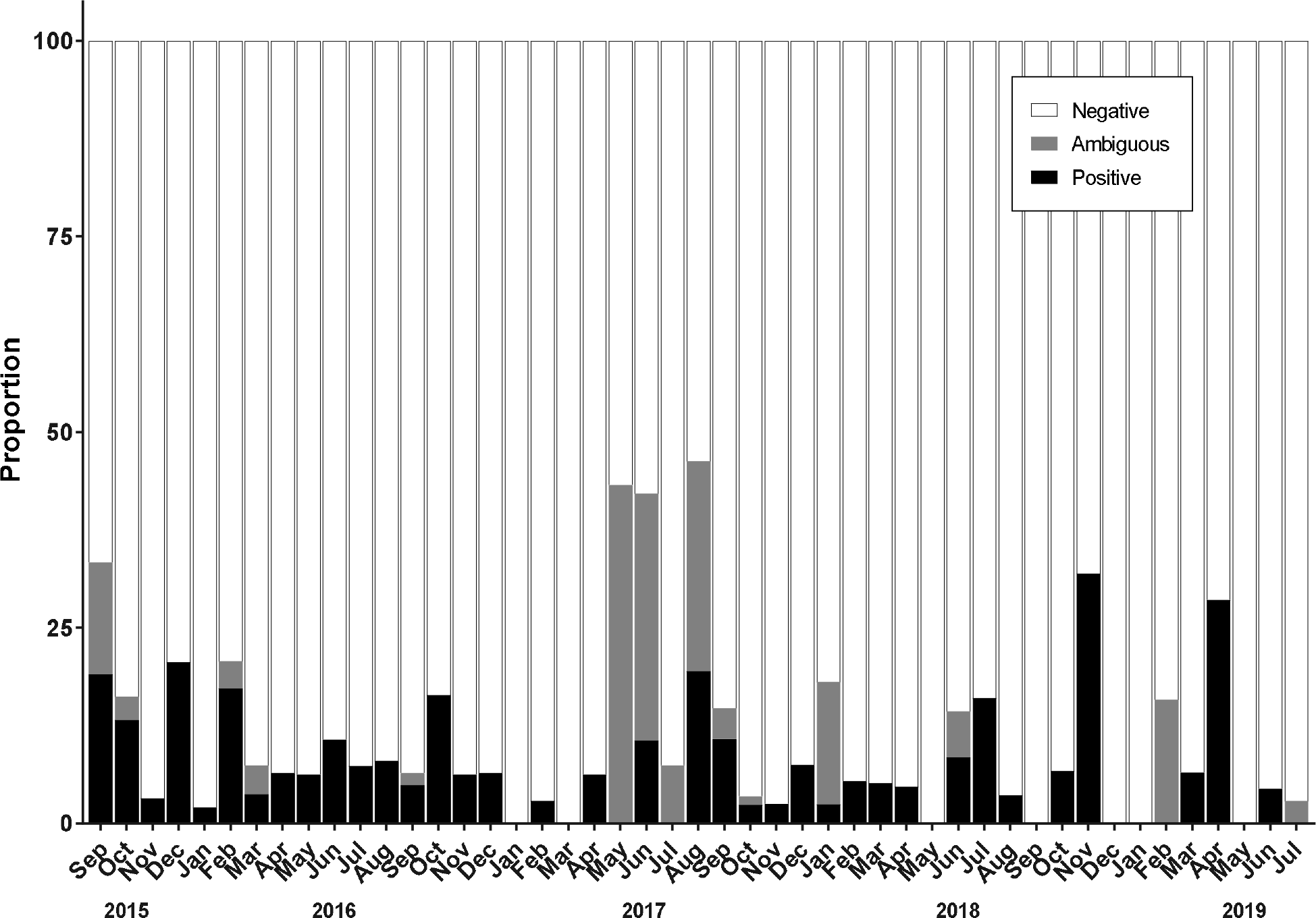
Mycoplasma testing results by month from September 2015 through July 2019.

For the final 12 months of data collection (beginning in July 2018), samples were further annotated with regard to their origin; ‘NCATS Internal’ or ‘Collaborator’. Any cell line in culture, or brought into culture from frozen stock after being cultured at NCATS was designated ‘NCATS Internal’. In a 12-month period, a larger proportion of collaborator samples (13%) than internal samples (5%) tested positive for mycoplasma. It is hoped that increasing awareness of the benefits of regular mycoplasma testing will reduce the number of positive cultures, particularly given that some cell lines provided from collaborator labs may have been utilized for research for many years unknown.

Occasionally, a mycoplasma contaminated cell line was treated with the anti-*Mycoplasma* compound plasmocin to remove the contamination^24^. Plasmocin is successful at removing contamination^24^ in most but not all cases, and studies with contaminated immortalized monocytes^25^ and human pluripotent stem cells demonstrated that plasmocin treatment did not alter cellular phenotypes (including stemness and pluripotency) compared with the original cell line^26^. Re-sourcing of cell lines is preferred, but not always an option. By way of example (anonymized to protect the guilty), NCATS scientists sourced a tumor cell line from the laboratory that created it. The cell line harbored a somatic mutation and was one of the few cell lines available to scientists at the time. However, on receiving the cell line, it was found to be highly contaminated and the originating lab was alerted. Discussions revealed that the lab had never tested for mycoplasma, had shared the cell line with many other labs (none of whom had raised the alarm), and that back-up stock was likely also contaminated. The NCATS scientists treated with plasmocin over multiple weeks, then cultured plasmocin-free for four weeks prior to testing and confirming that the cells were mycoplasma-free. The cell line was then shipped back to the originating lab as the new non-contaminated cell line stocks. By increasing awareness of the issue of mycoplasma contamination, we hope to encourage routine testing and provide suggestions for responding to contamination to lower the percentage of contaminated cell lines used in research laboratories.

### STR testing for cell line identity

In parallel, we established a routine practice for Short Tandem Repeat (STR) profiling for authentication of cell lines. STR profiling of human cell lines utilizes a PCR based assay to determine the number of repeated tandem DNA sequences at specific loci throughout the human genome^7^. The location of these sequences, or markers, is consistent for all humans, however each human has a differential number of short tandem repeated sequences at each marker, which were inherited parentally. By assessing multiple STR markers for each person, one can build a profile that can be compared to others to determine if the samples are from related or unrelated individuals. Standard methods for STR profiling involve using Promega kits which amplify 10, 16, or 18 markers per sample. Since each STR marker is present on both copies of each chromosome, each STR marker is reported as 2 numbers corresponding to the number of short tandem repeats at that marker.

If a cell line has a known STR profile (reference profile), then any sample purported to match it should be compared to the reference profile to ensure the sample is authentic. ICLAC standards state that related samples have greater than 80% of their STR alleles matching using the algorithm below:

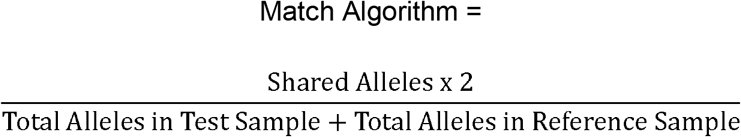

If a cell line does not have a known STR profile or is a newly generated cell line, it is important to establish the STR profile early on in order to track potential misidentification.

Frozen samples of cell lines were deposited in a common area with an associated request form specifying the number of STR markers to be profiled and the name of the cell line within the sample and passage number. A regular monthly shipment of samples were sent to a commercial provider of STR testing, primarily the Genetic Core Resources Facility at Johns Hopkins University STR profiling reports including the full profile as well as results of profile searches within ATCC and DSMZ databases were distributed to scientists. STR profiling is utilized by NCATS scientists to confirm the identity of human cell lines being used in our research laboratories, authenticate cell lines received from external collaborators (compared against reference STR profiles), or establish an STR profile for newly created cell lines.

To date, NCATS has tested 306 cell line samples for STR profiling, from 274 unique cell lines. Of these samples, 89 were primary and iPSC cell lines, where the goal was to ensure that subsequent clones had the same STR profiles as the original clone. Thirty samples had no published STR profile for reference, even though they were the cell lines were published in the literature. The remaining samples were cell lines with a known reference STR profile to compare against. Five cell lines did not match the known STR profile for that cell line (one example is described below), including a case where two cell lines being used by a scientist had been inadvertently switched. The remaining cell lines were >80% match (accepted threshold) to the available reference STR profile and were considered to be matches (Figure 3).

**Figure 2:**
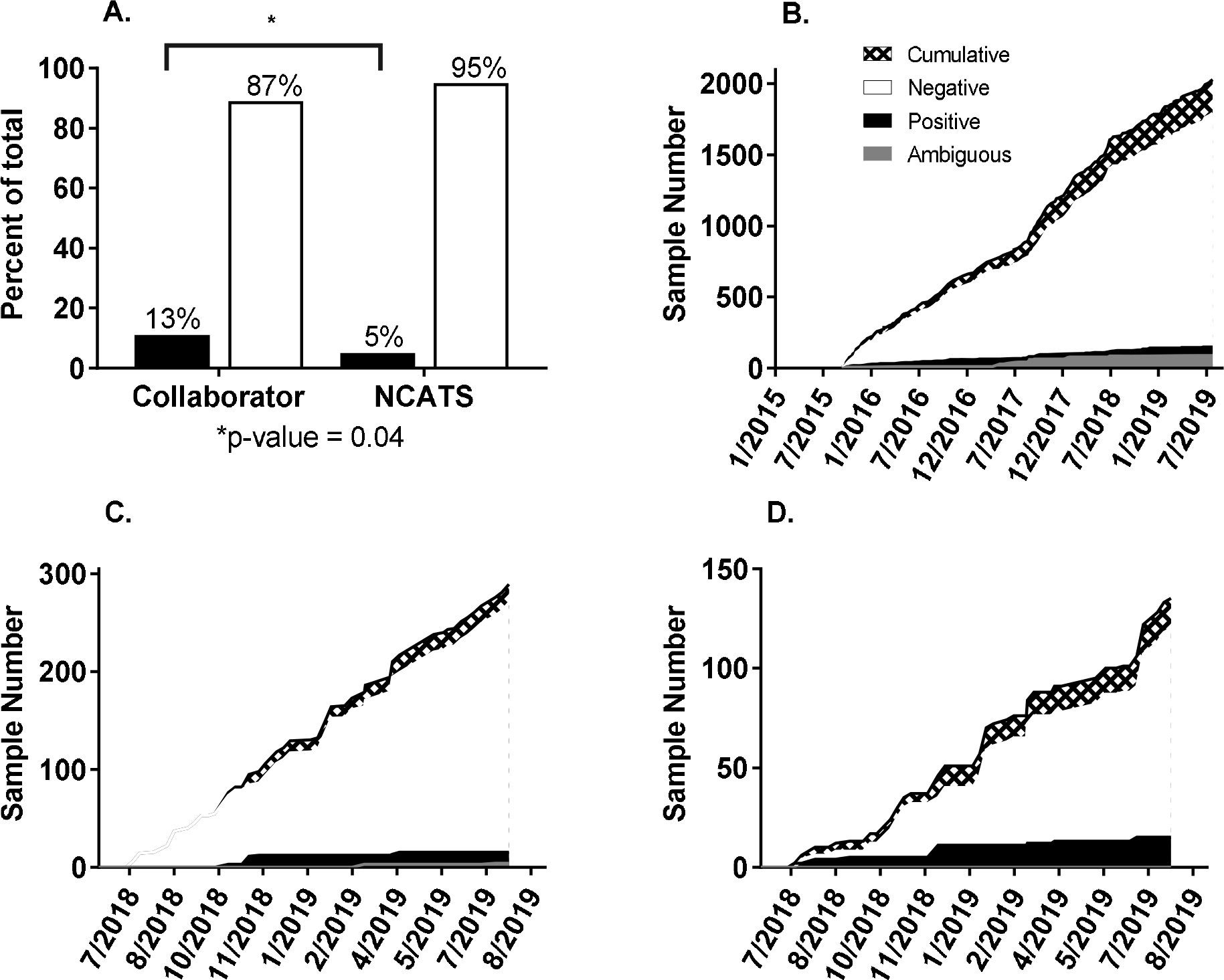
A) Retrospective analysis: Impact of sample origin on mycoplasma presence; n=135 and 289 for collaborator samples and NCATS samples, respectively. B) Cumulative results of all samples tested since July 2018. C) Cumulative results of NCATS samples. D) Cumulative results of collaborator samples.

**Figure 3:**
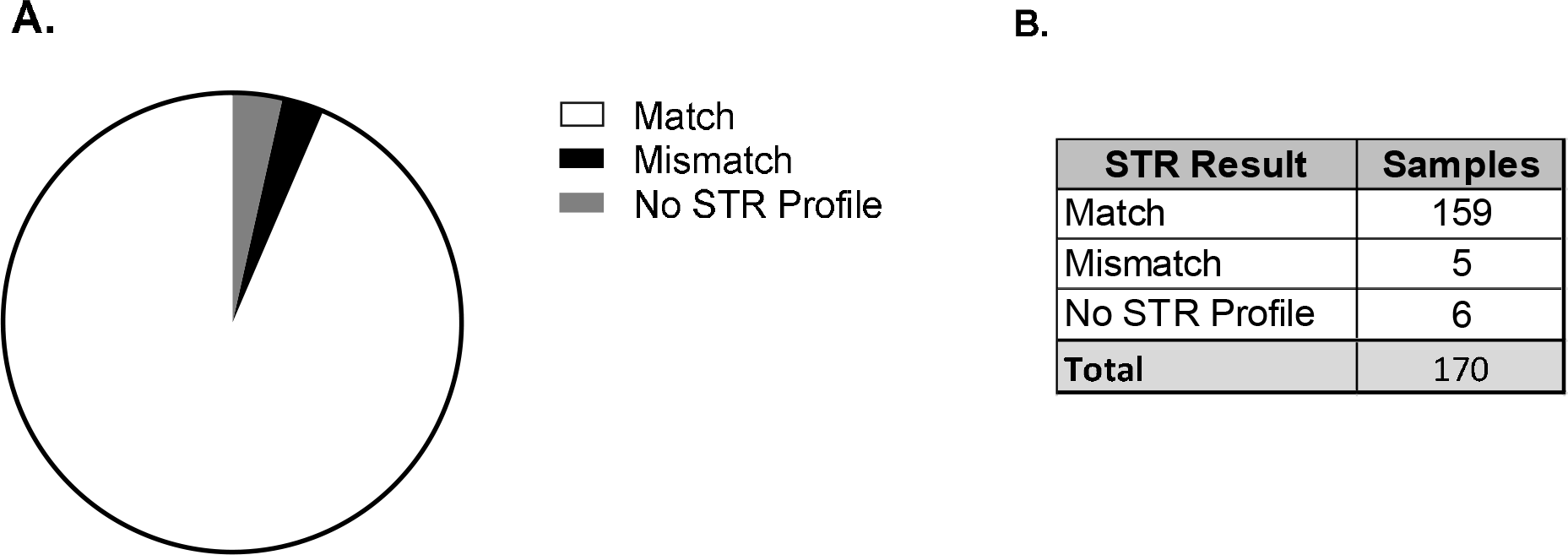
STR profiling results from 170 submitted samples; a >80% match to the available STR profile was accepted as a cell line match.

The benefit of a routine STR screening program can be made by the following example (as above, anonymized to protect the guilty). A collaboration was initiated with the goal of screening drug libraries against an isogenic pair of cell lines: a commonly used pancreatic cancer line BxPC3, and the same cell line with a stable knockdown of a specific gene. The aim was to identify approved drugs that may demonstrate synthetic lethal activity against the knockout line compared to wildtype, with the long-term goal of being able to take a validated active drug to the clinic. Upon receipt of the two cell lines, STR testing was performed, and it was found that the knockdown cell line was in fact another pancreatic cancer cell line (MiaPaCa-2). The collaborator had no earlier passage cells to utilize. A HTS using this mis-matched cell line pair could have been wasteful, with any bioactive hits from the screen leading to false conclusions regarding the translational potential of hits.

### Benefits of regular cell line testing

Given the high (>15%) tissue culture *Mycoplasma* contamination rates reported in the literature, it is likely that a significant number of published phenotypes may be affected by *Mycoplasma* contamination. At NCATS, systematic testing for *Mycoplasma* and STR profiles have prevented misidentified cell lines and contaminated cells from being used in high-throughput screening. This improves quality of research and the likelihood of reproducibility of our data.

To ensure that findings are robust, we suggest that phenotypes studied in long-term laboratory cell cultures be re-confirmed using the same cell line re-sourced from a cell line repository or originating laboratory.

